# Skin transcriptome reveals the Periodic changes in genes underlying hair follicle cycling in Cashmere goats

**DOI:** 10.1101/554030

**Authors:** Zhihong Liu, Feng Yang, Meng Zhao, Qing Mu, Tianyu Che, Yuchun Xie, Lina Ma, Lu Mi, Rui Su, Yanjun Zhang, Ruijun Wang, Zhiying Wang, Zhao Yanhong, Jinquan Li

**Affiliations:** College of Animal Science, Inner Mongolia Agricultural University, Hohhot, 010018, China.; Key Laboratory of Animal Genetics, Breeding and Reproduction, Inner Mongolia Autonomous Region, Hohhot, China.; Key Laboratory of Mutton Sheep Genetics and Breeding, Ministry of Agriculture, Hohhot, China.; Engineering Research Center for Goat Genetics and Breeding, Inner Mongolia Autonomous Region, Hohhot, China.

**Keywords:** Transcriptional group, differentially expressed genes, cashmere goat skin, villus growth cycle, keratin

## Abstract

Cashmere goats, as an important part of animal husbandry production, make outstanding contributions to animal fiber industry. In recent years, a great deal of research has been done on the molecular regulation mechanism of hair follicle cycle growth. However, there are few reports on the molecular regulation mechanisms of secondary hair follicle growth cycle in cashmere goats. In this study, we used transcriptome sequencing technique to sequence the skin of Inner Mongolia cashmere goats in different periods, Analyze the variation and difference of genes in the whole hair follicle cycle. And then, we verified the regulation mechanism of cashmere goat secondary hair follicle growth cycle by fluorescence quantitative PCR. As the result shows: The results of tissue section showed that the growth cycle of cashmere hair could be divided into three distinct periods: growth period (March-September), regression period (September-December) and resting period (December-March). The results of differential gene analysis showed that March was considered the beginning of the cycle, and the difference of gene expression was the most significant. Cluster analysis of gene expression in the whole growth cycle further supported the key nodes of the three periods of villus growth, and the differential gene expression of keratin corresponding to the villus growth cycle further supported the results of tissue slices. Quantitative fluorescence analysis showed that KAP3.1, KRTAP 8-1 and KRTAP 24-1 genes had close positive correlation with the growth cycle of cashmere, and their regulation was consistent with the growth cycle of cashmere. However, there was a sequence of expression time, indicating that the results of cycle regulation made the growth of cashmere change.

## Introduction

The growth of hair follicles in mammalian skin changes dynamically after birth, and the growth of hair follicles is cyclical. The hair growth cycle can be divided into the telogen, anagen, and catagen phases[1–5], each of which is regulated by specific genetic patterns[6, 7]. Inner Mongolia cashmere goats have two distinctly different fibrous structures, with thick and coarse hairs on the upper layer of the skin and fine and soft cashmere underneath. The cashmere comes from the secondary hair follicle structure in the skin[8], and the coarse hair comes from the primary hair follicle structure in the skin[9, 10]. Hair follicles, after shedding old hair shafts, produce new hair shafts[11], thereby starting a new cycle of hair follicle growth[12]. Keratin (KRT) and keratin-associated proteins (KRTAPs) are the main components of hair, affecting the physiological properties of hair. Hair follicle and hair shaft growth involve changes in the expression of genes encoding a KRT intermediate silk protein and a KRTAP.[13–16] Human hair follicles are not synchronized, and each hair follicle is independent of others[17]. In contrast, hair growth exhibits periodic changes with annual changes in daylight[17, 18]. This periodic growth pattern of hair depends on the intrinsic molecular mechanisms and external growth environment[19, 20].

With the progress of research technology, second-generation high-throughput sequencing technology can be used to screen for differentially expressed genes[21], allowing research across a broad spectrum of gene expression. RNA-Seq technology is a high-throughput sequencing technology that can be used for additional research on gene transcripts to discover low abundance transcripts and new transcripts and to identify differential expression of transcripts among different samples[22–24].

In this study, second-generation high-throughput sequencing technology was used for transcriptome sequencing of skin samples from different stages of hair growth. The aim of this study was to investigate the correlation between differentially expressed genes and the regulation of hair cycle transitions at different stages of hair growth. The biological functions of differentially expressed genes at different stages of hair growth play an important role in the elucidation of the regulatory mechanism of hair growth, laying a theoretical foundation for the study of this regulatory mechanism.

## Methods

### Animals

In this experiment, Three Inner Mongolian cashmere goats were selected from the same grazing environment. All animal experiments were performed in accordance with the ‘Guidelines for Experimental Animals’ of the Ministry of Science and Technology (Beijing, China). All surgery was performed according to recommendations proposed by the European Commission (1997), and was approved by experimental animal ethics committee of Inner Mongolia Agricultural University. These goats had the same weights, body sizes and ages; they were unrelated, and they exhibited good growth of does aged 1 year. The sampling period was 1 year, and none of the experimental goats were kids. Skin samples were collected at the beginning of each month. Skin samples were collected from the midline part of the scapula at 10-15 cm in the Department of Surgery, and directly stripped 3-cm-diameter skin samples were collected after flushing with PBS and rapid freezing in liquid nitrogen. Then, the samples were transferred to the laboratory in a liquid nitrogen tank and stored in a spare −80 °C freezer.

### Total RNA extraction from skin

Total RNA was extracted from the skin of three cashmere goats using an RNAiso Plus Kit (TRIzol method). The total RNA was tested for purity and integrity by a sterile UV-vis spectrophotometer and an Agilent 2100 bioanalyzer, respectively. Total RNA was stored in a freezer at −80 °C.

### Construction of sequencing library

CDNA library construction for transcriptome sequencing was performed according to the operating instructions for the Illumina TruSeqTM RNA Sample Preparation Kit. The total RNA from the three cashmere goats was pooled by mixing equal amounts of each. The mRNA was purified by using oligo-dT magnetic beads and then fragmented to 100-400-bp mRNA. Double-stranded cDNA was synthesized by using fragmented mRNA as a template, exonucleases and polymerases. The ends of the double-stranded cDNA fragments were blunted, and the double-stranded cDNAs were phosphorylated to ligate the sequencing adapters and poly (A) tail, and the sizes of the cDNA recovered were confirmed to be 200-300 bp by using a Bio-Rad Certified Low-Range Ultra Agarose Kit. PCR amplification of cDNA was performed to obtain a sequencing library, and library quality control was conducted using a TBS-380 instrument.

### Transcriptome sequencing

Paired-end sequencing of the cDNA was conducted by using an Illumina HiSeqTM 2000 sequencing platform. A 2 × 100 bp sequencing test was performed, and samples were sequenced by Shanghai Meiji Biopharmaceutical Co., Ltd.

### Quality and stitching of sequencing data and annotation of stitching results

The overlapping and low-quality sequencing data were removed. Trinity software was used to assemble the data into transcripts for transcriptional annotation and calculation of expression levels. The annotated sequences are aligned with the GO database using Blast2GO software, and annotated sequences were classified according to biological processes, cellular components and molecular functions. The expression levels were calculated using FPKM (fragments per kb of transcript per million mapped reads) [24], and finally, significant difference analysis was performed on the expression levels of all genes/transcripts in each group of samples. RSEM and R software were used to identify all the differentially expressed genes/transcripts, and R software was used to calculate the relative expression levels of the genes.

### Haematoxylin and eosin staining of skin samples over twelve months

Fresh skin samples were taken from Inner Mongolian cashmere goats over one year and fixed in 4% paraformaldehyde for 24 h, which was followed by dehydration in 30% sucrose solution for approximately 24 h. The samples were embedded in OCT, frozen and sectioned on a cryostat. The 6-μm-thick sliced tissue samples were strained with haematoxylin for one minute and rinsed with water. After washing, the sample was placed for 3 s in ethanol with 1% hydrochloric acid and then in 1% alkaline ethanol. Eosin staining was performed when the alcohol gradient was dehydrated to 95%, and dehydration was continued after staining. Then, the samples were treated with xylene and neutral gum and observed under a microscope.

### Quantitative reverse transcription PCR

The cDNA obtained above was used for quantitative reverse transcription PCR (q-PCR) analysis. The gene-specific primers used for q-PCR were designed with Primer 3.0 software and synthesized by Sangon Biotech Co., Ltd. (Shanghai, China). The primer sequences and fragment sizes are listed in Table 1. The q-PCR was performed using a PrimeScript RT Reagent Kit (TaKaRa) in a 20-μL reaction volume with 10 μL of 2× SYBR Premix Ex Taq II (TaKaRa), 2 μL of cDNA, and 0.5 μL of each primer. The reaction was assessed on a Bio-Rad IQ5 multicolour real-time PCR detection system (Hercules, CA, USA). The *β-actin* ware used as a reference. The qRT-PCR analysis was performed using the 2^−ΔΔCT^ method, and statistical analysis was performed by SPSS software (version 17.0). The values are represented as mean ± standard deviation. A significance level of 0.05 was used.

**Table 1.**
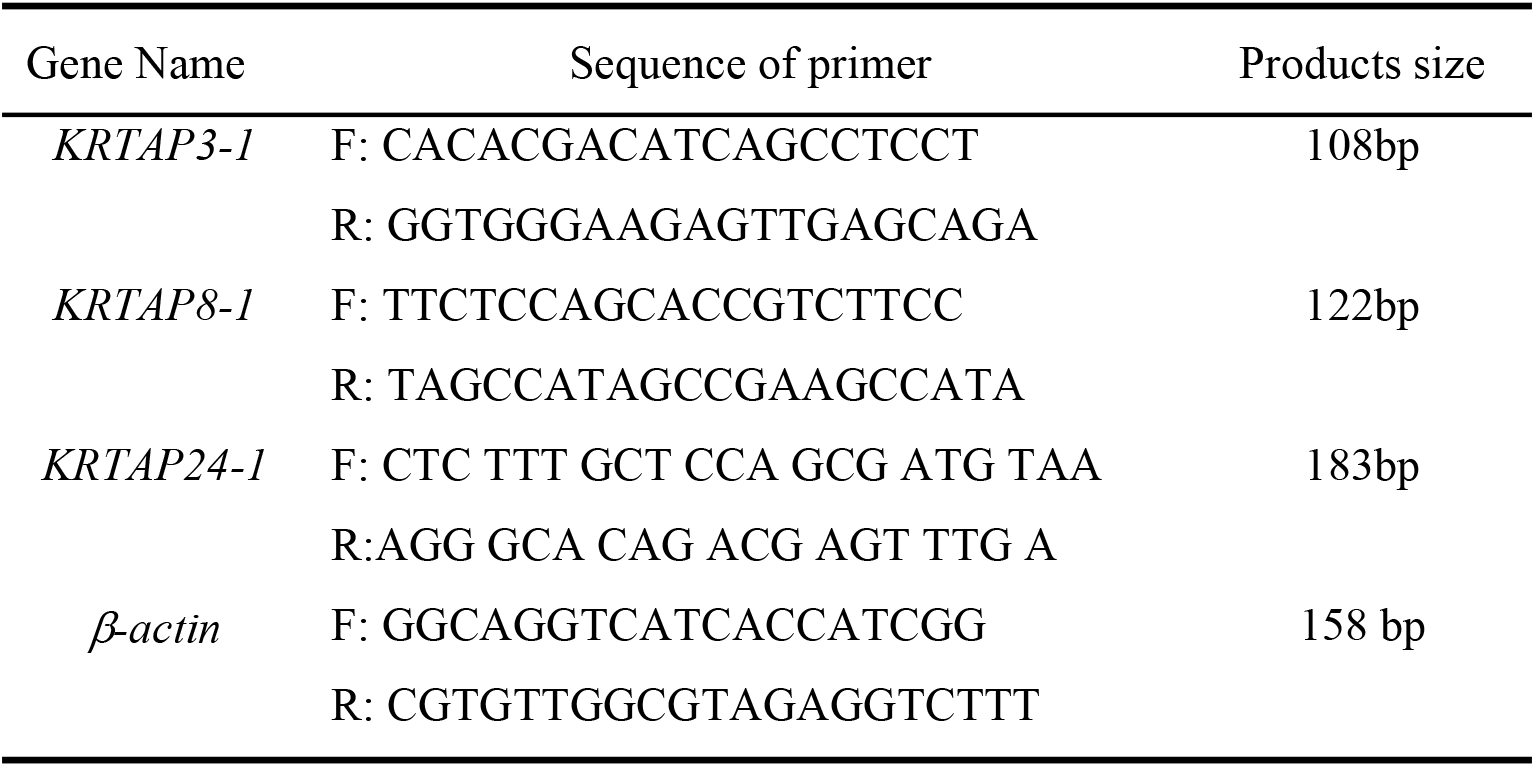
Primer sequences and fragment size of Cashmere goat *KRTAP3-1, KRTAP8-1, KRTAP24-1* gene and β-actin

## Results

### Morphological analysis of goat skin cycle changes

We observed the skin tissue of cashmere goats by tissue sectioning technology. As the results showed (Fig. 1), The number of secondary hair follicles in cashmere goats decreased gradually from December to March (Fig. 1 L, A, B and C). The lowest value was reached in March (Fig. 1 C), and the statistical values of each trait were also lower. The division and the extension to the dermis of hair follicles began in April (Fig. 1 D), The statistics of secondary hair follicles also began to increase at the same time. It is observed that there is a velvet appearance on July (Fig. 1 G). Most villi grow out of body surface from August to September (Fig. 1 H, I). At the same time, the statistical value of secondary follicles reached the highest level, This period is considered to be the peak period of villous growth. In October, hair follicle bulb cells began to enlarge, gradually aged and died, and the dermal papillae began to atrophy, The statistics of secondary hair follicles gradually decreased (Fig. 1 J). In December, the hair follicle roots rose to the sebaceous glands, and the secondary follicle statistics reached the lowest level (Fig. 1 L), This state has been maintained until February of next year. So we made the initial inference that The cycle of secondary hair follicles in cashmere goats can be divided into growth period from March to September, resting period from September to December and regression period from December to March.

**Fig. 1.**
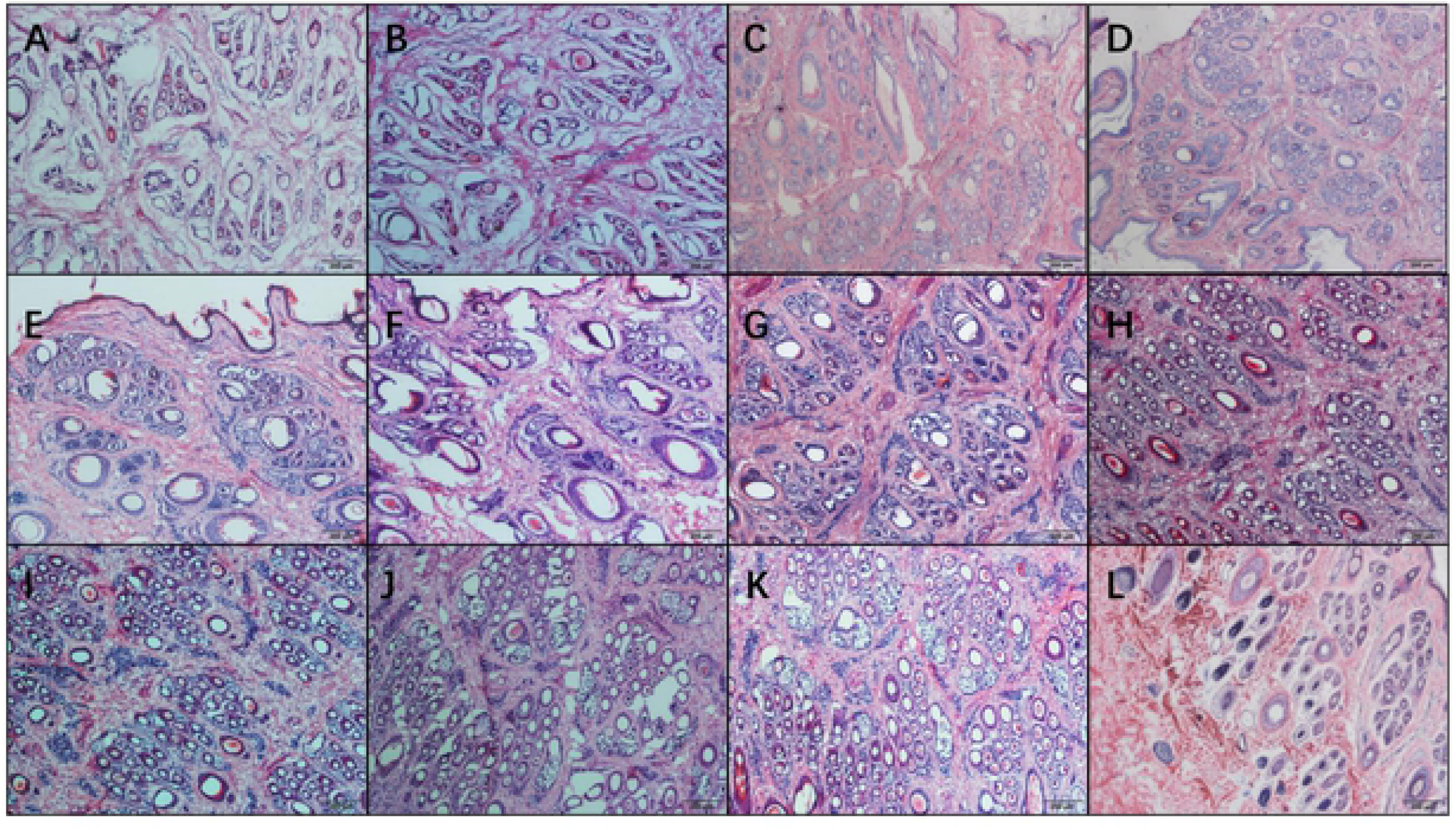
Morphological study on skin tissue of cashmere goat in 1-12 months. A January, B February, C March, D April, E May, F June, G July, H August, I September, J October, K November, L December

### Gene expression difference analysis

We first screened and analysed the data quality (Table 2) and data length distribution (Table 3).Then we compared the skin transcriptome data of cashmere goats in 12 months with each two seats (Fig. 2). It was found that the number of differentially expressed genes was the largest between February and March. The total number of differential genes was 1059, of which 219 were up-regulated and 840 were down regulated. In March and April, the number of differentially expressed genes was 731, of which 550 were up-regulated and 181 were down regulated. In June and July, there were 418 differentially expressed genes, of which 388 were up-regulated and 30 were down regulated. Those results showed that the expression of genes was initially up-regulated or down-regulated during the initiation of secondary hair follicle growth. Along with the advance of hair follicle initiation, the number of down-regulated genes began to decrease, and the number of up-regulated genes continued to increase. After the completion of the initiation process, the gene changes tended to be stable. The results further showed that the hair follicle development was initiated by the combination of up-regulation and down-regulation of genes in the early stage of initiation, and the gene expression returned to normal level after initiation. Comparing the data between June and July, we found that there was another significant change in gene expression during villus outgrowth. We believe that this change promotes villi to grow out of the body surface, but whether there are other roles remains to be further studied. From August to February of the next year, the secondary hair follicles genes expression changed significantly from quiescence to degeneration. This results further showed that the initiation of secondary hair follicles in cashmere goats began in March. Finally, it is worth mentioning that the number of differentially expressed genes increases first and then decreases from February to March and then to April, thus proving again that the secondary follicle cycle starts in March.

**Table 2.**
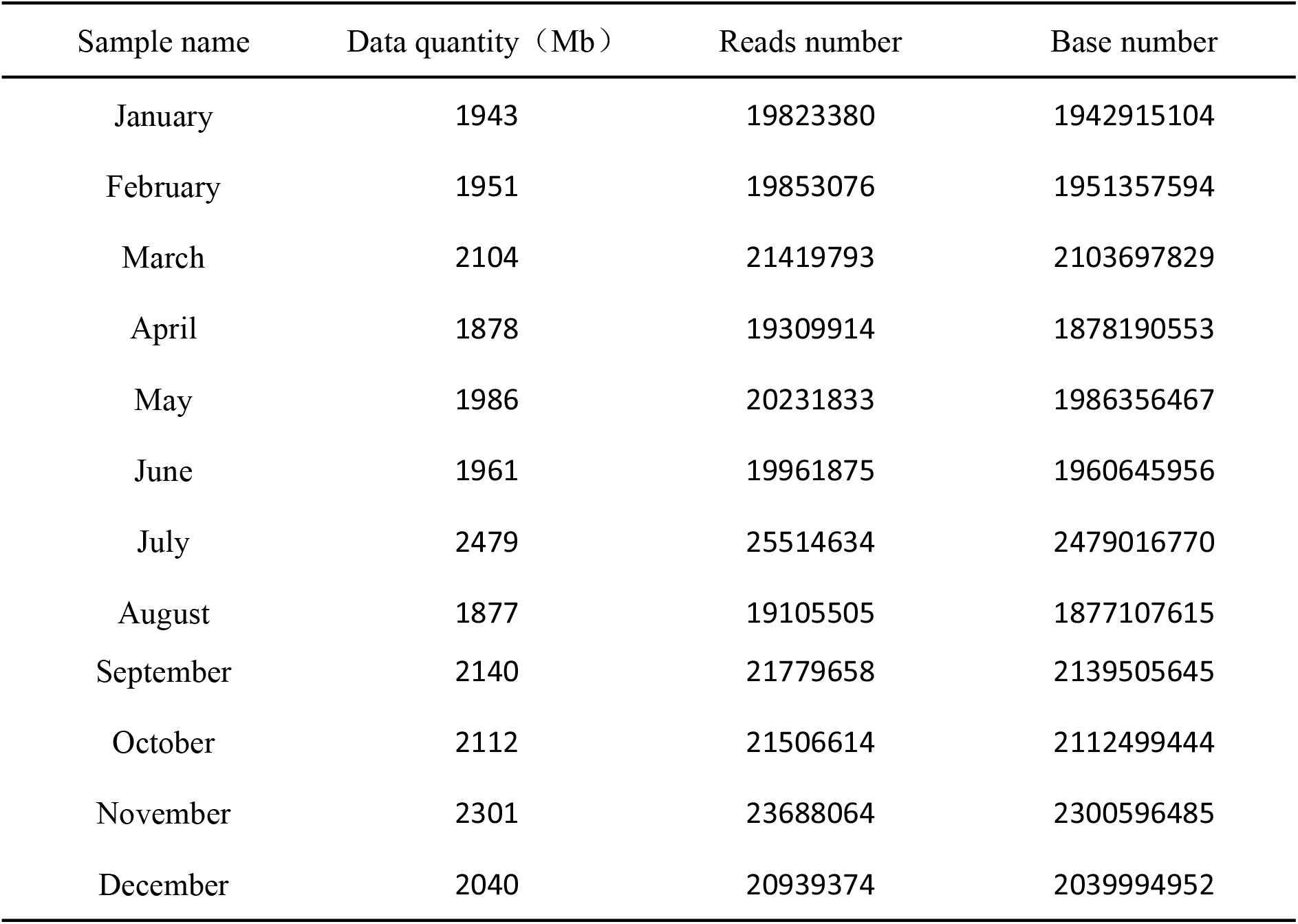
The result of the high quality raw data

| Sample name | Data quantity (Mb) | Reads number | Base number |
| --- | --- | --- | --- |
| January | 1943 | 19823380 | 1942915104 |
| February | 1951 | 19853076 | 1951357594 |
| March | 2104 | 21419793 | 2103697829 |
| April | 1878 | 19309914 | 1878190553 |
| May | 1986 | 20231833 | 1986356467 |
| June | 1961 | 19961875 | 1960645956 |
| July | 2479 | 25514634 | 2479016770 |
| August | 1877 | 19105505 | 1877107615 |
| September | 2140 | 21779658 | 2139505645 |
| October | 2112 | 21506614 | 2112499444 |
| November | 2301 | 23688064 | 2300596485 |
| December | 2040 | 20939374 | 2039994952 |

**Table 3.** Data of sequence length distribution

| Isogene length | Isogene number | Percentage (%) |
| --- | --- | --- |
| 1-400 | 7737 | 7.31 |
| 401-600 | 19281 | 18.21 |
| 601-800 | 11147 | 10.53 |
| 801-1000 | 7796 | 7.36 |
| 1001-1200 | 6054 | 5.72 |
| 1201-1400 | 5041 | 4.76 |
| 1401-1600 | 4545 | 4.29 |
| 1601-1800 | 4165 | 3.93 |
| 1801-2000 | 3755 | 3.54 |
| 2001-2200 | 3336 | 3.15 |
| 2201-2400 | 3076 | 2.91 |
| 2401-2600 | 2842 | 2.68 |
| 2601-2800 | 2624 | 2.48 |
| 2801-3000 | 2342 | 2.21 |
| 3001-3200 | 2106 | 2.00 |
| 3201-3400 | 1881 | 1.78 |
| 3401-3600 | 1802 | 1.70 |
| 3601-3800 | 1642 | 1.55 |
| 3801-4000 | 1430 | 1.35 |
| 4001-4200 | 1378 | 1.30 |
| 4201-4400 | 1182 | 1.12 |
| 4401-4600 | 1047 | 1.00 |
| ... | ... | ... |
| ALL | 105854 | 100 |

**Fig. 2.**
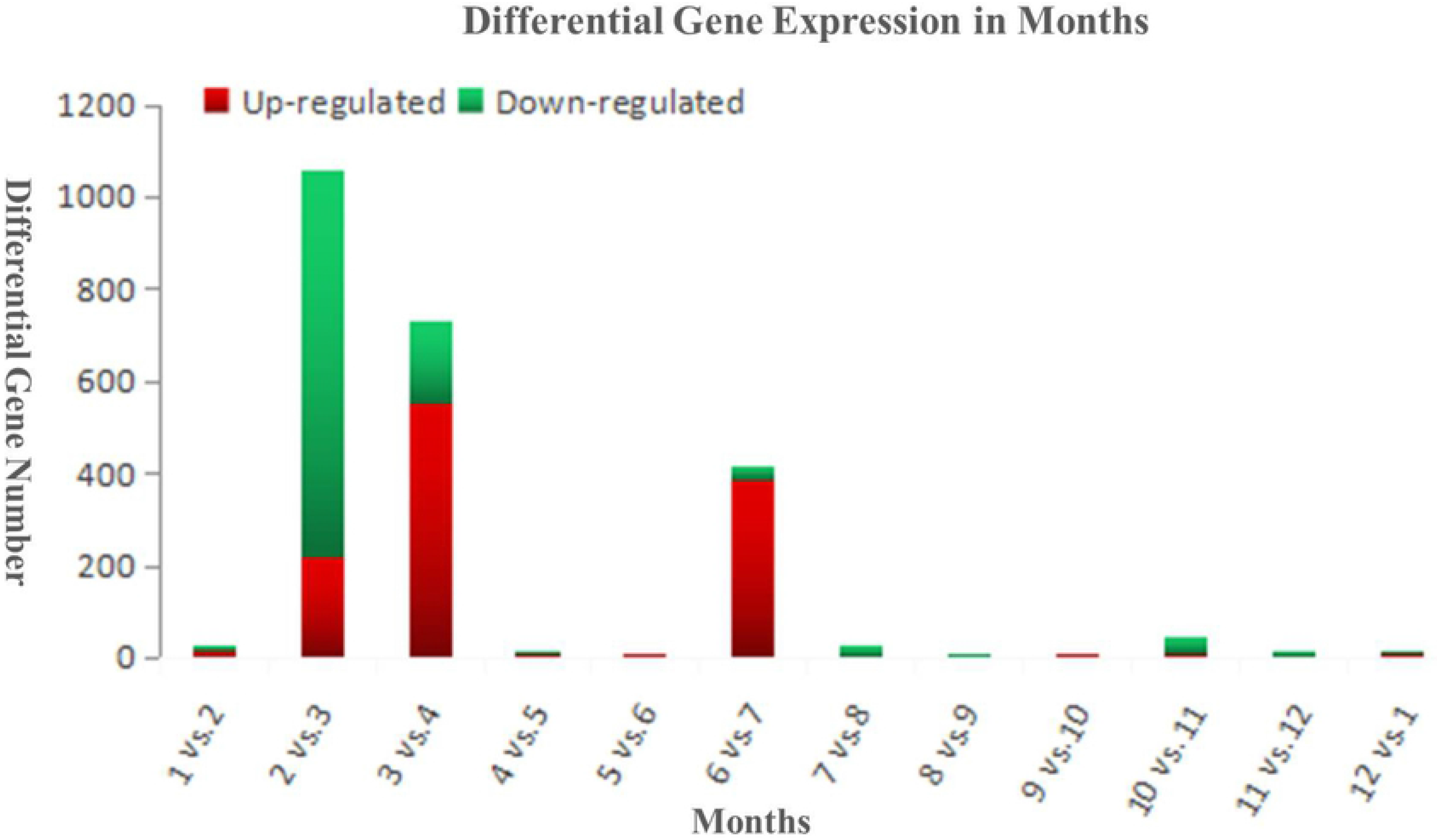
Histogram of differentially expressed gene statistics between neighbor month.

### Classification of gene function annotation

According to the GO classification statistics(Fig. 3), the skin expression genes can be divided into three main categories: biological functions, cell components and molecular functions. In this study, 51078 transcripts were noted with GO annotation. Among them, in biological function, the most annotated transcript is cellular process. In cellular component, most of the transcripts have been transcribed to cell and cell part. In molecular function, most of the transcripts have been transcribed to binding. It is speculated that during the hair follicle cycle, the changes of gene expression lead to the changes of the number and state of cells in the hair follicle structure, which further leads to the occurrence or shedding of villi.

**Fig. 3.**
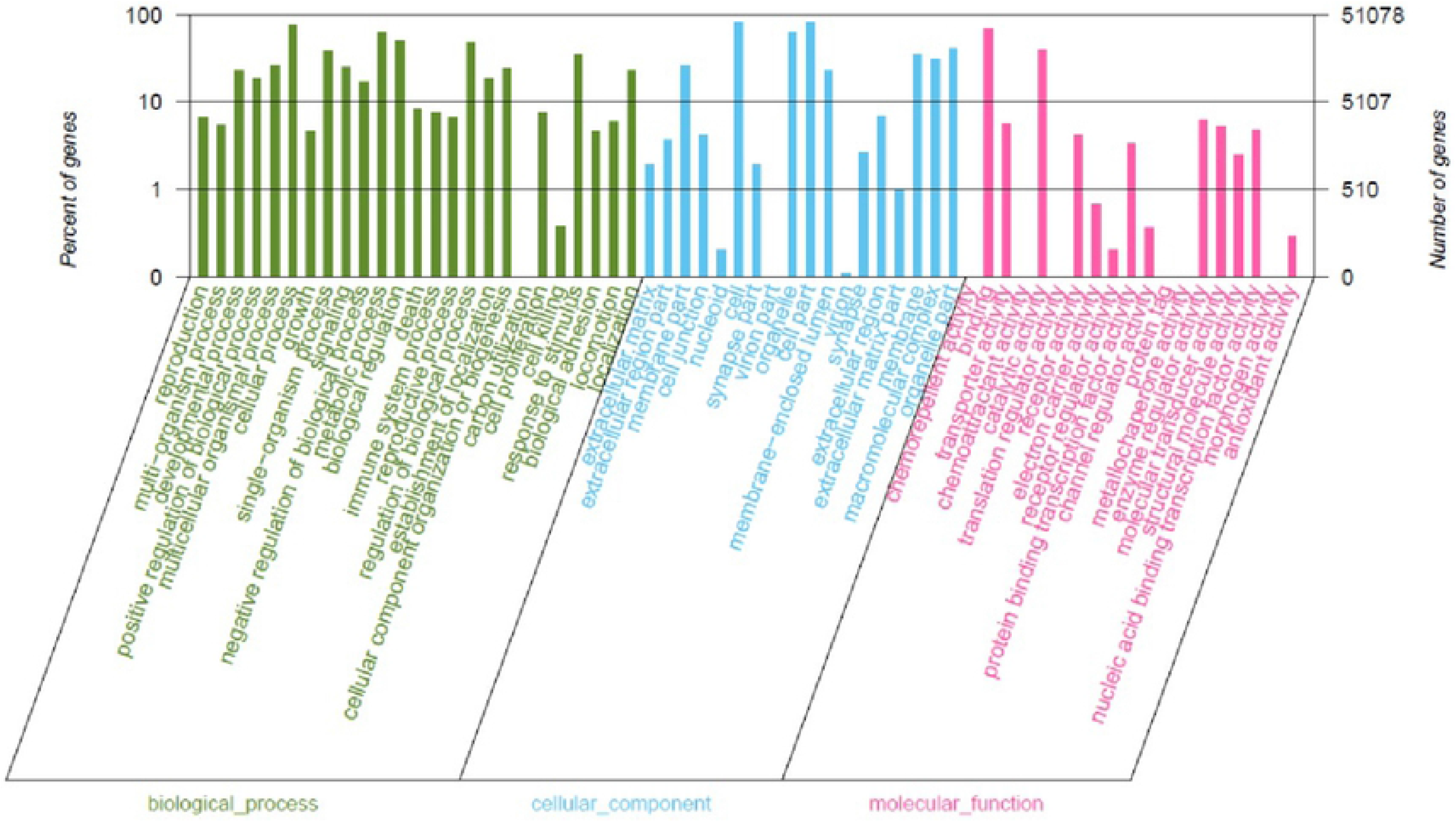
Gene ontology annotation.

### Group clustering analysis of natural periodic samples

In order to further explore the rule of gene expression, We calculated the correlation coefficient and cluster analysis of all gene expression levels in the 12 month natural cycle (Fig. 4). The results show that clustering information can be divided into three categories. The sample LZH3 was isolated because of the great changes in the gene expression of the follicle promoter. The sample LZH2-LZH7 ware considered to be the initiation process of hair follicle growth. Sample LZH8-LZH12 ware clustered together because it is thought to be the process of secondary hair follicles from vigorous growth to recession. Gene expression remained relatively unchanged from December to March, so the clustered sites were at the end of degeneration and before growth, They were considered to be the resting period of hair follicle development. Combined with previous studies in this article, we found that there ware several critical periods in the division of secondary hair follicle cycle. March is considered the key point for hair follicle cycle to start, September is the key period for the vigorous period of the hair follicle cycle and the beginning of the recession, December is the key point for the end of the hair follicle recession and the beginning of the rest period, These three critical periods determine the key signals in hair follicle and villi growth.

**Fig. 4.**
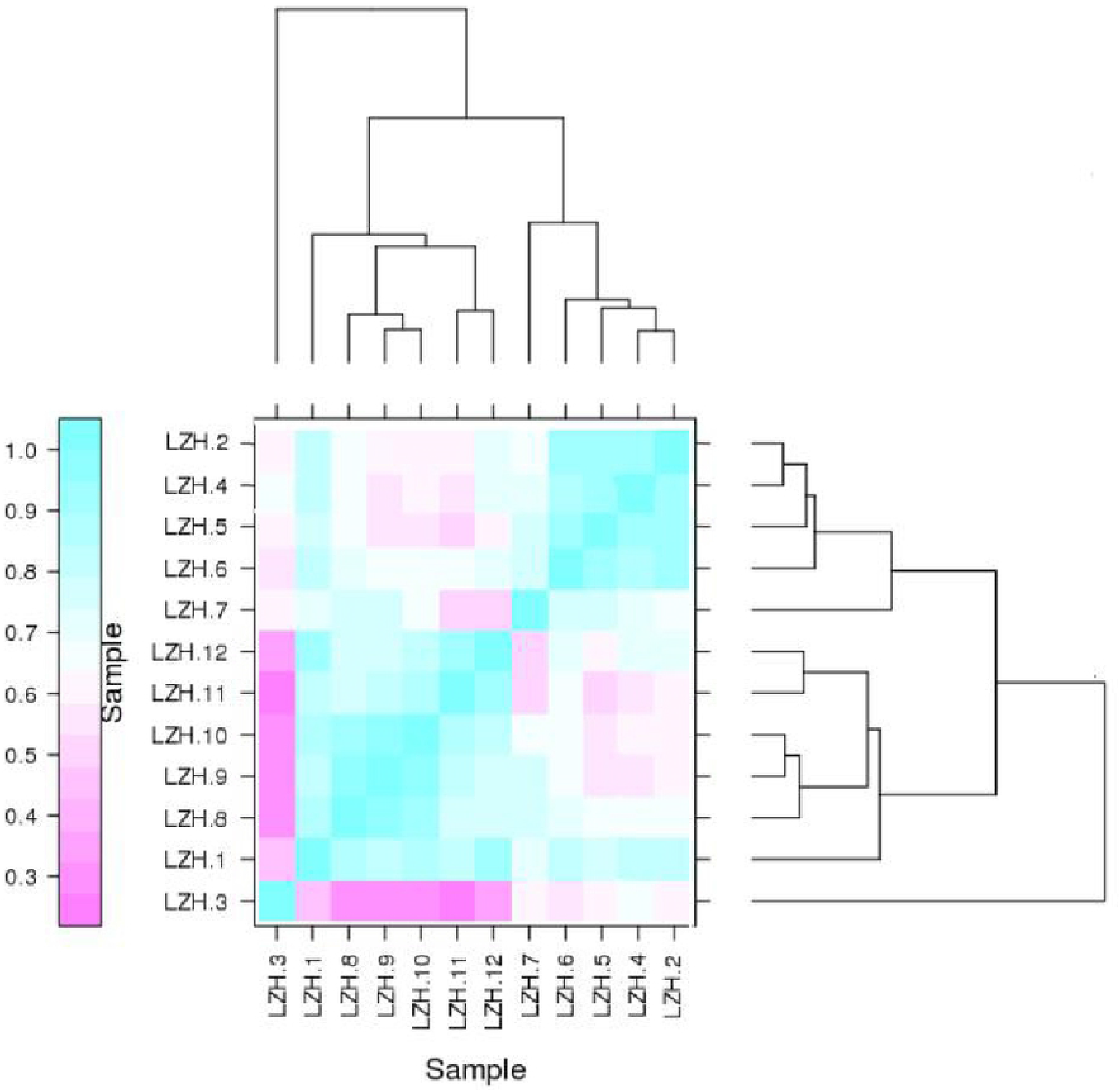
Cluster diagram of the growth cycle of cashmere.

### Extraction and analysis of target gene expression information

In order to explore the expression patterns of genes that play a key role in the cycle, we extract the expression information of all target genes for 1-12 months, and then cluster the expression patterns by analysis and exclusion. Gene expression patterns of several pathways related to villus cycle were obtained (Fig. 5A). The results showed that the expression pattern of genes related to villus cycle was consistent with our analysis of differential gene expression. The results further supported that villus of villus cycle began to enter the start stage in March, entered the regression stage in September and entered the end stage in December. However, the development process of villus cycle can not be visualized through the skin appearance results, because the development of cycle precedes the growth of villus, and there is a causal relationship between them. The period of villus growth is visually observed by tissue sections, and there is a direct relationship between villus growth and the expression of keratin. Therefore, in order to further verify the villus growth cycle, we also clustered the expression patterns of keratin and keratin-related genes (Fig. 5B). The results showed that the expression of keratin was consistent with the results of tissue sections, which further supported that the villus cycle first started (degenerated or rested), and then cascaded leading to changes in the expression of keratin gene, thus promoting the occurrence of villus (growth or degeneration). In order to further study the relationship between keratin gene and cycle, we take the gene with the highest expression as a case for further study.

**Fig. 5.**
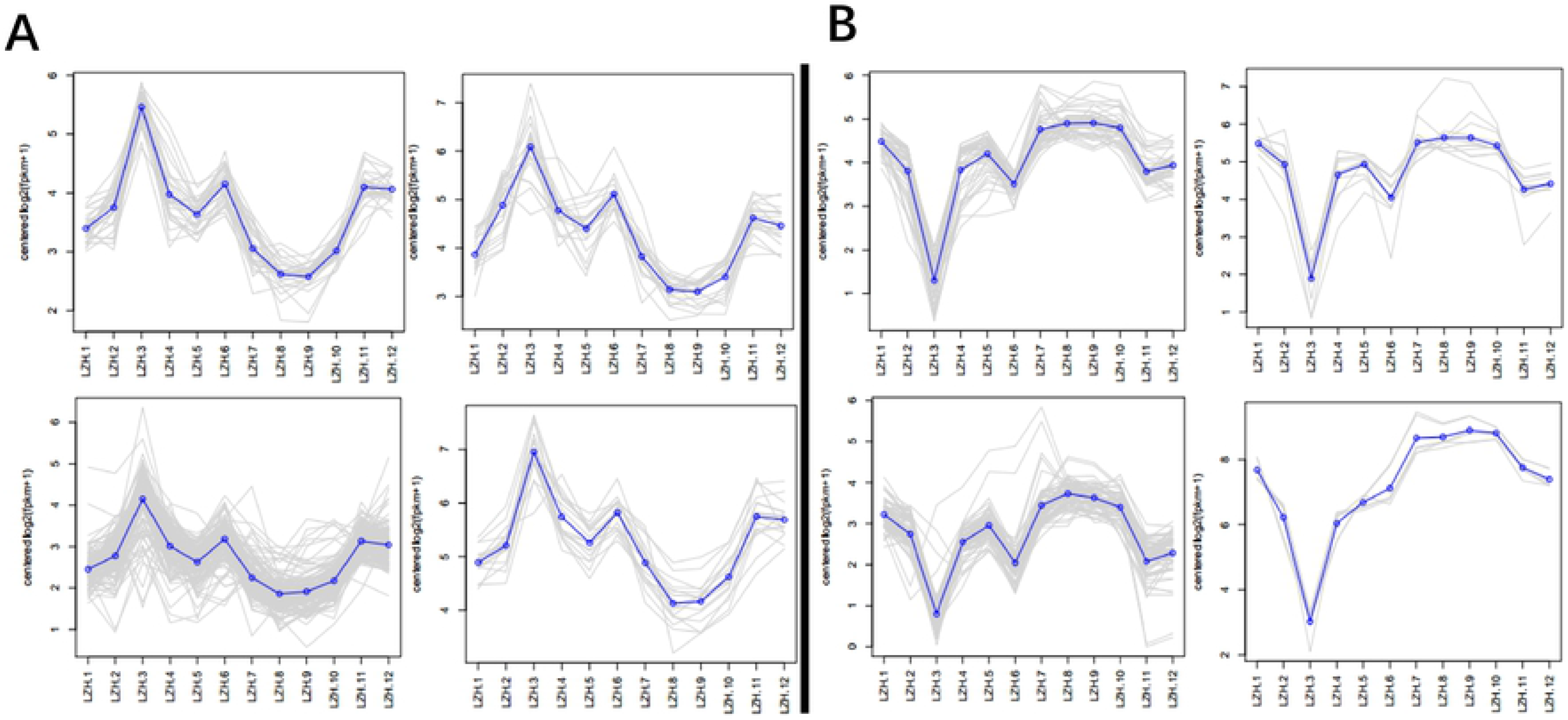
Clustered the expression patterns.

### QPCR analysis of key genes

It is well known that during the development of cashmere goat hair follicles, the changes of gene expression make the expression of keratin change and promote the development of villi. In order to further verify the role of genes in hair follicle cycle, we detected the changes of expression of several important keratins in cashmere goats for 12 months, and reversely verified the role of gene changes in cycle regulation. The expression of keratin associated protein 3-1 in cashmere for 12 months ware confirmed by quantitative PCR (Fig. 6 A). The results showed that *KAP 3-1* was expressed in the skin at different stages of a year, but the expression of *KAP 3-1* was significantly different (P<0.05) and fluctuated periodically in 12 months. The expression level of three months in August, September and October was significantly higher than that in other months (P < 0.05). Combined with previous studies, the expression quantity was verified by dividing different periods (Fig. 6 B). It was found that the expression of *KAP3-1* gene in growth phase was significantly higher than that in rest or regression phase. Subsequently, To verify the stability of gene expression. We examined the expression of two other genes *KAP 8-1* and *KAP 24-1* in cashmere goat skin at several stages by fluorescence quantitative analysis (Fig. 7). The relative expression of *KRTAP 8-1* (Fig. 7A) and *KRTAP 24-1* (Fig. 7B) genes in Inner Mongolia cashmere goats’ skin showed periodic variation, which was consistent with the hair follicle development cycle. It is indicated that *KRTAP 8-1* and *KRTAP 24-1* genes played a positive role in controlling cashmere wool growth and closely related to the regulation of villi growth and cycle transformation.

**Fig. 6.**
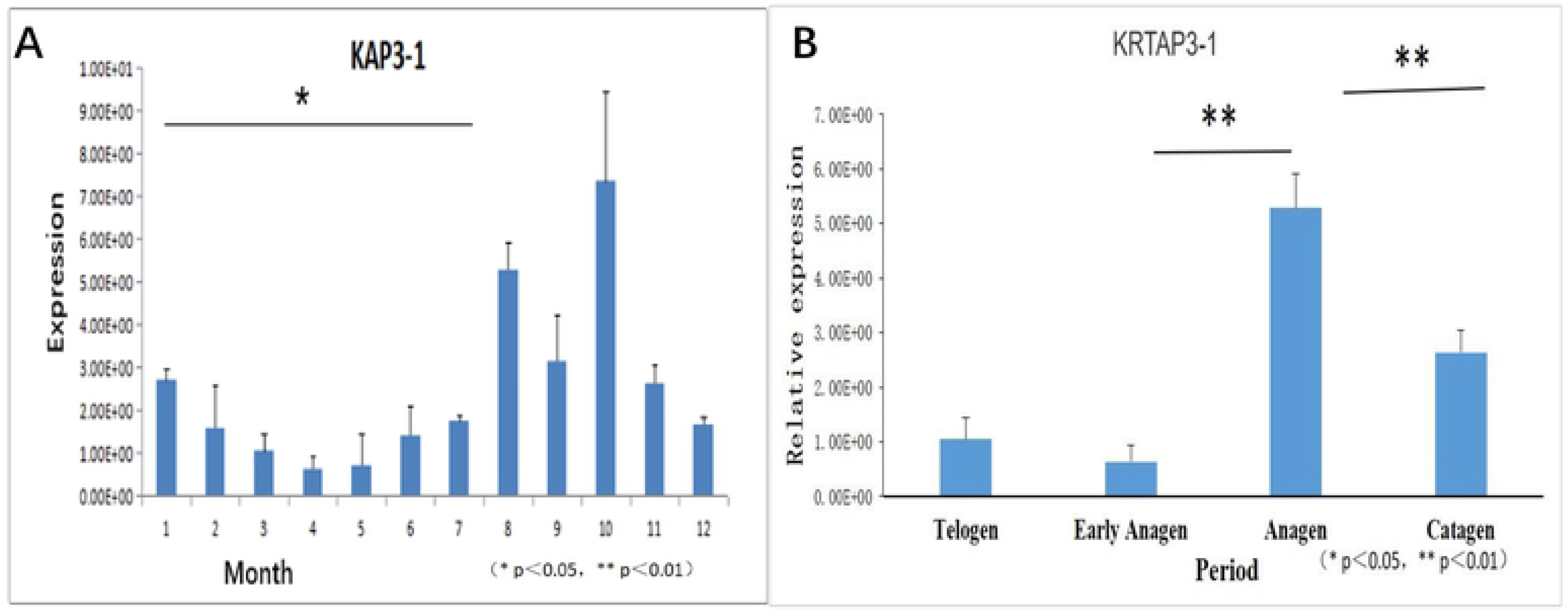
KRTAP 3-1 change in the amounts of expression in different month and periods.

**Fig. 7.**
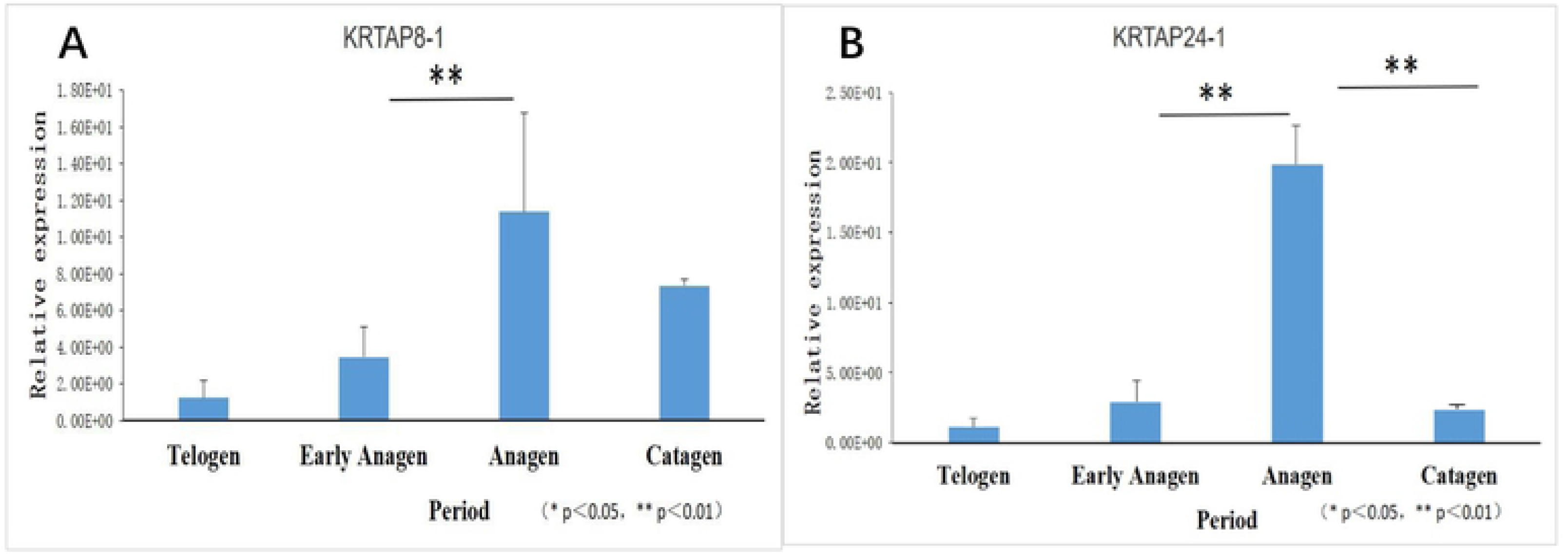
KRTAP 8-1 and KRTAP 24-1 change in the amounts of expression in different periods.

## Discussion

In natural environment, there is a regular growth pattern of animal hair. The growth of animal hair follicles is constantly changing when birthed[25, 26]. The hair follicles have the characteristics of self-renewal and periodic growth. Hair follicle growth is periodic, which can be divided into anagen, catagen and telogen[27–29]. Therefore, it is important to study the changes of cashmere goat hair follicles and the expression differences of their regulatory genes to improve the economic value of cashmere goats.

Secondary hair follicle development of cashmere goats is a cyclical process[30]. In this study, the histological slices of cashmere goat secondary follicles showed that the skin thickness, the length and depth of primary follicles, the width, density and activity of primary follicles did not change significantly from January to March, and the statistical value of each trait was low. From April onwards, the cell division in the root of hair follicle accelerated and extended to the dermis, and the morphological data began to increase until July, when the villi grew out of the body surface. In August and September, most of the villus grew out of the body surface. At this time, most of the statistical values of the characteristics of the follicle reached the annual maximum, which was the peak period of villus growth. From October, hair follicle globular cells began to senile and die, and hair papilla cells began to atrophy. At this time, the statistical value of hair follicle morphological data began to decline and become smaller. In December, the root of hair follicle rose to the vicinity of sebaceous gland, and the statistical value of hair follicle reached the lowest level in the whole year until February of the following year. It is inferred that the growth period of cashmere goat hair follicles is from March to September, the regression period is from September to December, and the rest period is from December to March. The results are consistent with previous preliminary studies[20, 31–33].

Through transcriptome sequencing technology, we can find out the changing rules of skin expression genes in cashmere goats at different stages and the expression situation at different stages[21, 34, 35]. Transcriptome sequencing can guide us to study the direction of hair follicle cycle in gene expression. However, we observed the phenotypic characteristics of hair follicles by tissue sections, and preliminarily explored the expression regularity of hair follicle cycle and the differences of hair follicle characteristics in different periods. Then, transcriptome sequencing was used to detect gene expression and expression pattern, and the expression level and expression pattern were obtained to verify the results of tissue sections. Finally, the results of gene expression were verified by fluorescent RT-qPCR. This reverse validation is used to ensure the accuracy of phenotypic traits and gene expression patterns. More accurately locating and studying genes related to hair follicle growth cycle. The results of this study showed that the number of secondary hair follicles began to increase from March. The results of transcriptome sequencing showed that there were significant changes in gene expression in March compared with February, and the gene expression increased, and the number of differentially expressed genes increased. These two results verify each other and infer that March is the start of the secondary hair follicle cycle. In September, the villus grew to its peak, then the villus began to decrease, and the number of secondary follicles began to decrease. Meanwhile, Gene expression was first down-regulated and then up-regulated in the three months from August to October, and gene changes were obvious in September, which indicated that September was the end of villus growth cycle and the beginning of degeneration. The number of hair follicles in degenerative period lasted until December, and then the number of secondary hair follicles remained unchanged from December to early March of next year, that is, the period of the end of the villus growth cycle.

KRT and KRTAPs contribute nearly 90% of the cashmere yield[36–38], demonstrating indirectly that the composition and interactions of these proteins play an important role in cashmere quality[39–41]. KRT and KRTAP gene expression directly affects cashmere fineness and density and other characteristics[42]. Most of the genes identified by transcript sequencing of Inner Mongolian cashmere goat were members of the KRT and KRTAP gene families, which indicated that the expression of KRTAPs directly affected the growth of hair follicles and cashmere-related traits.

Thus, the cycle of hair follicle growth is correlated with variation in gene expression, and the complex regulates the cycle of hair follicle growth[43]. In addition, there is a sharp fluctuation in the expression of genes during the telogen and anagen phases. However, genetic variation in the growth of cashmere stops growth relatively slowly, which is a gradual process. Therefore, the study of genes related to the initiation of cashmere growth has potential value in the discovery of genes that affect cashmere growth in cashmere goats, such as genes that regulate the cashmere cycle and the growth of cashmere, and the changes in expression from paused growth to active growth are more severe at the genetic level, which is obviously useful for gene expression research.

## Conclusion and Summary

In this study, we used transcriptome sequencing technique to sequence the skin of Inner Mongolia cashmere goats in different periods, Analyze the variation and difference of genes in the whole hair follicle cycle. And then, we verified the regulation mechanism of cashmere goat secondary hair follicle growth cycle by fluorescence quantitative PCR. As the result shows: The results of tissue section showed that the growth cycle of cashmere hair could be divided into three distinct periods: growth period (March-September), regression period (September-December) and resting period (December-March). The results of differential gene analysis showed that March was considered the beginning of the cycle, and the difference of gene expression was the most significant. Cluster analysis of gene expression in the whole growth cycle further supported the key nodes of the three periods of villus growth, and the differential gene expression of keratin corresponding to the villus growth cycle further supported the results of tissue slices. Quantitative fluorescence analysis showed that KAP3.1, KRTAP 8-1 and KRTAP 24-1 genes had close positive correlation with the growth cycle of cashmere, and their regulation was consistent with the growth cycle of cashmere. However, there was a sequence of expression time, indicating that the results of cycle regulation made the growth of cashmere change.

## Acknowledgements

The authors are thankful for the samples provided by the Aerbasi White Cashmere Goat Breeding Farm. Yanhong Zhao and Jinquan Li provided the test platform, whereas as co-first author of this article, Zhihong Liu and Feng Yang helped in designing and conducting the experiments and the analysis, evaluation, and interpretation of the results.

## Author contributions

FY, QM,YC and MZ made substantial contributions to the conception and design of the experiments. Conceived and designed the experiments:FY, ZL, JL. Performed the experiments: FY,ZL, QM, MZ, HZ. Analyzed the data: MZ, FY, ZL. Wrote the paper: RS, FY,HL. Critically revised the manuscript: FY, ZL,JL. All authors read and approved the final manuscript.

## References

1. Wu Z, Sun L, Liu G et al. Hair follicle development and related gene and protein expression of skins in Rex rabbits during the first 8 weeks of life. Asian-Australasian journal of animal sciences doi:10.5713/ajas.18.0256 (2018).

2. Millar SE. Molecular mechanisms regulating hair follicle development. The Journal of investigative dermatology 118(2), 216–225 (2002).

3. Messenger AG, Botchkareva NV. Unraveling the secret life of the hair follicle: from fungi to innovative hair loss therapies. Experimental dermatology 26(6), 471 (2017).

4. Botchkarev VA, Paus R. Molecular biology of hair morphogenesis: development and cycling. Journal of experimental zoology. Part B, Molecular and developmental evolution 298(1), 164–180 (2003).

5. Watabe R, Yamaguchi T, Kabashima-Kubo R, Yoshioka M, Nishio D, Nakamura M. Leptin controls hair follicle cycling. Experimental dermatology 23(4), 228–229 (2014).

6. Fuchs E. Scratching the surface of skin development. Nature 445(7130), 834–842 (2007).

7. Mikkola ML. Genetic basis of skin appendage development. Seminars in cell & developmental biology 18(2), 225–236 (2007).

8. Zhu B, Xu T, Yuan J, Guo X, Liu D. Transcriptome sequencing reveals differences between primary and secondary hair follicle-derived dermal papilla cells of the Cashmere goat (Capra hircus). PloS one 8(9), e76282 (2013).

9. Bykov YS, Schaffer M, Dodonova SO et al. The structure of the COPI coat determined within the cell. eLife 6 (2017).

10. Pin D, Cachon T, Carozzo C. Determination of the depth of excision using a dermatome (Aesculap) to export all hair follicle bulbs from a donor site in the dog. Journal of veterinary medicine. A, Physiology, pathology, clinical medicine 54(9), 539–541 (2007).

11. Ahmad SA, Way M. New Editor on Journal of Cell Science. Journal of cell science 130(2), 303 (2017).

12. Sennett R, Rendl M. Mesenchymal-epithelial interactions during hair follicle morphogenesis and cycling. Seminars in cell & developmental biology 23(8), 917–927 (2012).

13. Shimomura Y, Aoki N, Rogers MA, Langbein L, Schweizer J, Ito M. hKAP1.6 and hKAP1.7, two novel human high sulfur keratin-associated proteins are expressed in the hair follicle cortex. The Journal of investigative dermatology 118(2), 226–231 (2002).

14. Rogers MA, Langbein L, Praetzel-Wunder S, Giehl K. Characterization and expression analysis of the hair keratin associated protein KAP26.1. The British journal of dermatology 159(3), 725–729 (2008).

15. Rogers MA, Langbein L, Winter H, Ehmann C, Praetzel S, Schweizer J. Characterization of a first domain of human high glycine-tyrosine and high sulfur keratin-associated protein (KAP) genes on chromosome 21q22.1. The Journal of biological chemistry 277(50), 48993–49002 (2002).

16. Pruett ND, Tkatchenko TV, Jave-Suarez L et al. Krtap16, characterization of a new hair keratin-associated protein (KAP) gene complex on mouse chromosome 16 and evidence for regulation by Hoxc13. The Journal of biological chemistry 279(49), 51524–51533 (2004).

17. Plikus MV, Baker RE, Chen CC et al. Self-organizing and stochastic behaviors during the regeneration of hair stem cells. Science 332(6029), 586–589 (2011).

18. Fu S, Zhao H, Zheng Z, Li J, Zhang W. Melatonin regulating the expression of miRNAs involved in hair follicle cycle of cashmere goats skin. Yi chuan = Hereditas 36(12), 1235–1242 (2014).

19. Paus R, Burgoa I, Platt CI, Griffiths T, Poblet E, Izeta A. Biology of the eyelash hair follicle: an enigma in plain sight. The British journal of dermatology 174(4), 741–752 (2016).

20. Yang M, Song S, Dong K et al. Skin transcriptome reveals the intrinsic molecular mechanisms underlying hair follicle cycling in Cashmere goats under natural and shortened photoperiod conditions. Sci Rep 7(1), 13502 (2017).

21. Wang L, Cai B, Zhou S et al. RNA-seq reveals transcriptome changes in goats following myostatin gene knockout. PloS one 12(12), e0187966 (2017).

22. Wang Z, Gerstein M, Snyder M. RNA-Seq: a revolutionary tool for transcriptomics. Nature reviews. Genetics 10(1), 57–63 (2009).

23. Nakamura T, Yabuta Y, Okamoto I et al. SC3-seq: a method for highly parallel and quantitative measurement of single-cell gene expression. Nucleic acids research 43(9), e60 (2015).

24. Ozsolak F, Milos PM. RNA sequencing: advances, challenges and opportunities. Nature reviews. Genetics 12(2), 87–98 (2011).

25. Cetera M, Leybova L, Woo FW, Deans M, Devenport D. Planar cell polarity-dependent and independent functions in the emergence of tissue-scale hair follicle patterns. Developmental biology 428(1), 188–203 (2017).

26. Balana ME, Charreau HE, Leiros GJ. Epidermal stem cells and skin tissue engineering in hair follicle regeneration. World journal of stem cells 7(4), 711–727 (2015).

27. Geyfman M, Plikus MV, Treffeisen E, Andersen B, Paus R. Resting no more: re-defining telogen, the maintenance stage of the hair growth cycle. Biological reviews of the Cambridge Philosophical Society 90(4), 1179–1196 (2015).

28. Hu HM, Zhang SB, Lei XH et al. Estrogen leads to reversible hair cycle retardation through inducing premature catagen and maintaining telogen. PloS one 7(7), e40124 (2012).

29. Ahmed NS, Ghatak S, El Masry MS et al. Epidermal E-Cadherin Dependent beta-Catenin Pathway Is Phytochemical Inducible and Accelerates Anagen Hair Cycling. Molecular therapy : the journal of the American Society of Gene Therapy 25(11), 2502–2512 (2017).

30. Rile N, Liu Z, Gao L et al. Expression of Vimentin in hair follicle growth cycle of inner Mongolian Cashmere goats. BMC genomics 19(1), 38 (2018).

31. Geng R, Yuan C, Chen Y. Exploring differentially expressed genes by RNA-Seq in cashmere goat (Capra hircus) skin during hair follicle development and cycling. PloS one 8(4), e62704 (2013).

32. Han W, Li X, Wang L et al. Expression of fox-related genes in the skin follicles of Inner Mongolia cashmere goat. Asian-Australasian journal of animal sciences 31(3), 316–326 (2018).

33. Wang S, Ge W, Luo Z et al. Integrated analysis of coding genes and non-coding RNAs during hair follicle cycle of cashmere goat (Capra hircus). BMC genomics 18(1), 767 (2017).

34. Wang Q, Oh JW, Lee HL et al. A multi-scale model for hair follicles reveals heterogeneous domains driving rapid spatiotemporal hair growth patterning. eLife 6 (2017).

35. Fb UB, Cau L, Tafazzoli A et al. Mutations in Three Genes Encoding Proteins Involved in Hair Shaft Formation Cause Uncombable Hair Syndrome. American journal of human genetics 99(6), 1292–1304 (2016).

36. Jin M, Wang J, Chu MX, Piao J, Piao JA, Zhao FQ. The Study on Biological Function of Keratin 26, a Novel Member of Liaoning Cashmere Goat Keratin Gene Family. PloS one 11(12), e0168015 (2016).

37. Wang F, Zieman A, Coulombe PA. Skin Keratins. Methods in enzymology 568 303–350 (2016).

38. Wang S, Luo Z, Zhang Y, Yuan D, Ge W, Wang X. The inconsistent regulation of HOXC13 on different keratins and the regulation mechanism on HOXC13 in cashmere goat (Capra hircus). BMC genomics 19(1), 630 (2018).

39. Birbeck MS, Mercer EH, Barnicot NA. The structure and formation of pigment granules in human hair. Experimental cell research 10(2), 505–514 (1956).

40. Yuan S, Li F, Meng Q et al. Post-transcriptional Regulation of Keratinocyte Progenitor Cell Expansion, Differentiation and Hair Follicle Regression by miR-22. PLoS genetics 11(5), e1005253 (2015).

41. Parry DA, Strelkov SV, Burkhard P, Aebi U, Herrmann H. Towards a molecular description of intermediate filament structure and assembly. Experimental cell research 313(10), 2204–2216 (2007).

42. Khan I, Maldonado E, Vasconcelos V, O’brien SJ, Johnson WE, Antunes A. Mammalian keratin associated proteins (KRTAPs) subgenomes: disentangling hair diversity and adaptation to terrestrial and aquatic environments. BMC genomics 15 779 (2014).

43. Rahmatalla SA, Arends D, Reissmann M et al. Whole genome population genetics analysis of Sudanese goats identifies regions harboring genes associated with major traits. BMC genetics 18(1), 92 (2017).

